# Achieving Resilience in Aging: How Mitochondrial Modulation Drives Age-associated Fluconazole Tolerance in *Cryptococcus neoformans*

**DOI:** 10.1101/2024.03.26.586817

**Authors:** Kyungyoon Yoo, Natalia Kronbauer Oliveira, Somanon Bhattacharya, Bettina C. Fries

**Affiliations:** Department of Microbiology and Immunology, Renaissance School of Medicine, Stony Brook University, Stony Brook, NY 11794, USA; Division of Infectious Diseases, Department of Medicine, Stony Brook University, Stony Brook, NY 11794, USA; Veterans Administration Medical Center, Northport, NY 11768, USA

**Keywords:** *Cryptococcus neoformans*, antifungal tolerance, replicative aging, mitochondria, reactive oxygen species, ergosterol, ABC transporter

## Abstract

*Cryptococcus neoformans* (*Cn*) is an opportunistic fungal microorganism that causes life-threatening meningoencephalitis. During the infection, the microbial population is heterogeneously composed of cells with varying generational ages, with older cells accumulating during chronic infections. This is attributed to their enhanced resistance to phagocytic killing and tolerance of antifungals like fluconazole (FLC). In this study, we investigated the role of ergosterol synthesis, ATP-binding cassette (ABC) transporters, and mitochondrial metabolism in the regulation of age-dependent FLC tolerance. We find that old *Cn* cells increase the production of ergosterol and exhibit upregulation of ABC transporters. Old cells also show transcriptional and phenotypic characteristics consistent with increased metabolic activity, leading to increased ATP production. This is accompanied by increased production of reactive oxygen species (ROS), which results in mitochondrial fragmentation. This study demonstrates that the metabolic changes occurring in the mitochondria of old cells drive the increase in ergosterol synthesis and the upregulation of ABC transporters, leading to FLC tolerance.

**IMPORTANCE:** Infections caused by *Cryptococcus neoformans* cause more than 180,000 deaths annually. Estimated one-year mortality for patients receiving care ranges from 20% in developed countries to 70% in developing countries, suggesting that current treatments are inadequate. Some fungal cells can persist and replicate despite the usage of current antifungal regimens, leading to death or treatment failure. In replicative aging, older cells display a resilient phenotype, characterized by their enhanced tolerance against antifungals and resistance to killing by host cells. This study shows that age-dependent increase in mitochondrial reactive oxygen species drive changes in ABC transporters and ergosterol synthesis, ultimately leading to the heightened tolerance against fluconazole in old *C. neoformans* cells. Understanding the underlying molecular mechanisms of this age-associated antifungal tolerance will enable more targeted antifungal therapies for cryptococcal infections.

## INTRODUCTION

*Cryptococcus neoformans* (*Cn*) is an opportunistic fungal pathogen that most commonly infects patients with advanced HIV infection (1–3), and it is responsible for 19% of AIDS-related deaths (4). The most serious manifestation of cryptococcosis is chronic meningoencephalitis (CME), which is often fatal. Estimated one-year mortality for patients receiving care ranges from 20% in developed countries to 70% in developing countries (4), suggesting that current treatments are inadequate (5).

In countries with limited resources where CME is most prevalent, fluconazole (FLC) is often the only available antifungal used for induction, consolidation, and long-term maintenance regimes (6). FLC works by inhibiting 14-alpha-demethylase, one of the key enzymes in ergosterol synthesis, thereby targeting an important component of the fungal plasma membrane (7).

One factor that contributes to the failure of FLC-based therapy is the development of drug tolerance, or the ability of a subpopulation of cells to survive and grow in the presence of the drug at drug concentrations above the minimum inhibitory concentration (MIC) (8–11). As opposed to FLC resistance, FLC tolerance is not identified by standard MIC assays. FLC resistance in *Cn* can be mediated by changes in the ergosterol target and augmentation of drug efflux pumps, where mutations in *ERG* genes of the ergosterol biosynthesis pathway decrease the susceptibility to FLC (12, 13), and overexpression of efflux pumps decreases intracellular drug concentration (14, 15). The efflux of FLC is facilitated by the ATP-binding cassette (ABC) transporters, which require ATP to drive transport out of the cell (14). Most of the ATP synthesis occurs in mitochondria, and biosynthesis of ergosterol starts from acetyl-coenzyme A (acetyl-CoA), a key mitochondrial intermediate. Furthermore, altered mitochondrial regulation has also been linked to FLC resistance in several pathogenic fungi (16–18). However, the connection among ergosterol regulation, ABC transporters, and mitochondrial metabolism and how it contributes to FLC tolerance is unknown.

Like other pathogenic fungi, *Cn* undergoes asymmetric cell division during replicative aging, where age is measured by the number of generations (19–21). During this process, proteins are unequally distributed between the mother and daughter cells, leading to age-dependent phenotypes (22). Older *Cn* cells that have undergone aging are more resistant to antifungals (15, 23) and macrophage killing (24). As a result, older cells accumulate in cerebrospinal fluid both in a rat model and in humans during infection (25), supporting the concept that replicative aging conveys enhanced resilience within the host environment.

Cells of advanced generational age constitute the dominant subpopulation *in vivo*, which can be attributed to their enhanced FLC tolerance (25, 26). However, the underlying mechanism behind the age-dependent FLC tolerance remains unknown. In this study, we explore the interplay between ergosterol regulation, mitochondrial metabolism, and ABC transporters in mediating age-dependent FLC tolerance in old *Cn* cells. We show that old *Cn* cells exhibit altered metabolic activity, leading to increased ATP production that fuels ABC transporters, as well as upregulation of ergosterol synthesis. We further demonstrate that the increased mitochondrial burden in old cells leads to ROS-mediated mitochondrial signaling, resulting in alteration in ergosterol and ABC transporter dynamics that contribute to age-associated FLC tolerance. These results offer insight into the molecular dynamics governing age-associated mitochondrial signaling with its repercussions on drug tolerance, providing a foundation for more targeted therapeutic interventions.

## RESULTS

### Transcriptomic Impact of Replicative Aging in *Cryptococcus neoformans*

To investigate the effects of replicative aging on gene expression in *Cn*, we performed RNA-seq analyses on young and old *Cn* cells. Aging resulted in the differential regulation of 1266 genes (adjusted p-value ≤ 0.05), including 725 upregulated and 541 downregulated genes (Fig 1A). Heat map visualization of RNA-seq data and Pearson’s Correlation Analysis of the biological replicates indicated the reliability of our RNA-seq results (Fig S1). Further exploration using gene ontology (GO) enrichment analysis unveiled an upregulation of genes associated with 27 biological processes (BPs) (Fig 1B). Additionally, genes related to 19 molecular functions (MFs) were upregulated in old cells (Fig 1C). Conversely, aging correlated with the downregulation of genes linked to 34 biological processes and 23 molecular functions (Fig S2).

**Figure 1.**
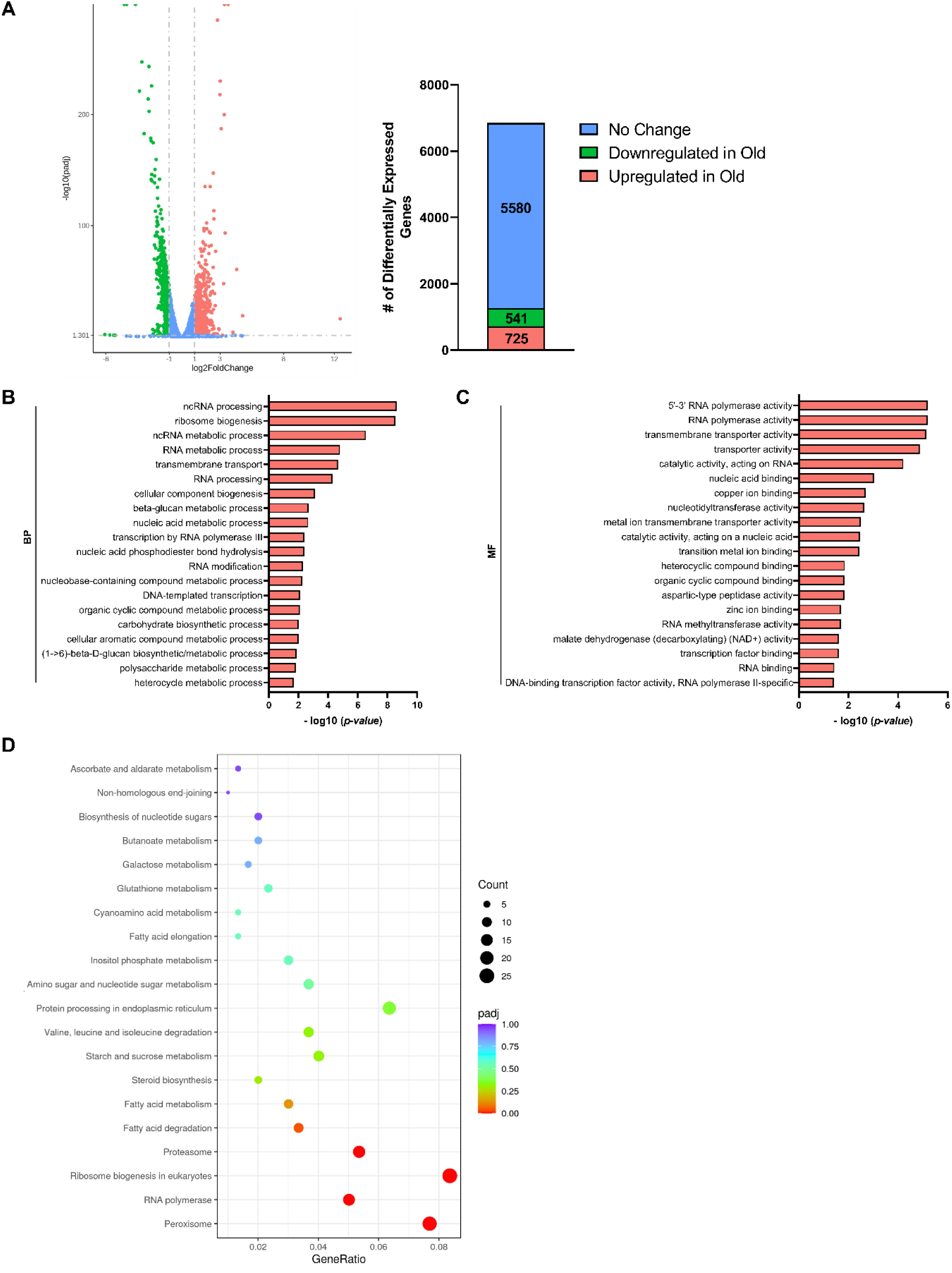
Replicative Aging Results in Transcriptomic Changes Related to Metabolic Processes in *C. neoformans*. RNA-seq of young (1-3 generations) and old (10 generations) *Cn* cells was performed. (A) Volcano plot of genes that are differentially regulated in old and young cells. Genes that are upregulated in old vs. young (Log_2_FC ≥ 1, adjusted p ≤ 0.05) are indicated by pink points while genes that are downregulated in old vs. young (Log_2_FC ≤ −1, adjusted p ≤ 0.05) are indicated by green points. Genes that do not meet these criteria are shown in blue. Gene ontology (GO) terms of differentially expressed genes show (B) molecular function (MF) and (C) biological process (BP) that are upregulated in old *Cn* cells. Presented are the top twenty GO terms based on the lowest over-represented p-values. according (D) Kyoto Encyclopedia of Genes and Genomes (KEGG) analysis of differentially expressed genes shows molecular pathways that are upregulated in old *Cn* cells.

Kyoto Encyclopedia of Genes and Genomes (KEGG) pathway analysis also indicated that some important pathways, such as peroxisome, fatty acid degradation, and steroid biosynthesis, were enriched in old *Cn* cells (Fig 1D). One of the key functions of peroxisomes is the β-oxidation of fatty acids (27), which produce acetyl-CoA that can serve as a substrate for ATP production through the tricarboxylic acid (TCA) cycle. Furthermore, acetyl-CoA is the central metabolite for the biosynthesis of ergosterol, the most abundant sterol in fungal cell membranes. Because ATP-powered ABC transporters and ergosterol biosynthesis are both implicated in mediating FLC tolerance, we looked more closely at how aging affects gene expression of these aspects of metabolism. As indicated by the KEGG analysis, various genes that encode enzymes in sterol biosynthesis and fatty acid oxidation were upregulated in old *Cn* cells (Table 1). Genes associated with respiratory functions, such as ones encoding cytochrome C oxidases II and III, apocytochrome C, and NADH dehydrogenases. Furthermore, genes encoding various mitochondrial functions also exhibited significant upregulation in aged cells. Altogether, these results from the transcriptomic analysis of young and old *Cn* cells indicate that aging affects various metabolic pathways that are linked with FLC tolerance.

**Table 1.** Subset of genes upregulated in old *Cn* cells.

| Function | Gene | Function | Log<sub>2</sub>FC<br>(Old/Young) | P-Adj |
| --- | --- | --- | --- | --- |
| Sterol<br>Biosynthesis | CKF44_01737 | Methylsterol monooxygenase | 1.16 | 1.76E-16 |
|  | CKF44_03819 | Sterol 24-C-methyltransferase | 1.49 | 6.26E-48 |
|  | CKF44_00519 | lathosterol oxidase | 1.27 | 2.19E-25 |
|  | CKF44_00040 | sterol 14-demethylase | 1.11 | 8.62E-24 |
|  | CKF44_06829 | squalene monooxygenase | 1.16 | 2.50E-17 |
| Fatty Acid<br>Oxidation | CKF44_06010 | Aldehyde dehydrogenase (NAD) | 1.11 | 3.17E-28 |
|  | CKF44_04553 | 3-hydroxyacyl-CoA dehydrogenase | 1.16 | 3.41E-05 |
|  | CKF44_04640 | ATP-citrate synthase subunit 1 | 1.18 | 8.44E-51 |
| Respiration | CKF44_09012 | Cytochrome c oxidase subunit II | 1.12 | 2.08E-07 |
|  | CKF44_09004 | Cytochrome c oxidase subunit III | 1.12 | 0.0050 |
|  | CKF44_09002 | Apocytochrome b | 1.95 | 0.025 |
|  | CKF44_00149 | NADH dehydrogenase<br>(ubiquinone) 1 alpha subcomplex 4 | 1.17 | 3.80E-22 |
|  | CKF44_00788 | NADH dehydrogenase,<br>mitochondrial | 1.74 | 1.40E-56 |
| Mitochondrial<br>Function | CKF44_07409 | Mitochondrial DNA-directed RNA<br>polymerase | 1.19 | 3.85E-16 |
|  | CKF44_01288 | Mitochondrial carrier protein | 1.83 | 1.05E-10 |
|  | CKF44_00861 | 3-hydroxyacyl-CoA dehydrogenase | 1.95 | 7.47E-05 |
|  | CKF44_00268 | Mitochondrial acetolactate<br>synthase | 1.08 | 3.27E-18 |

### Integration of Transcriptomic and Phenotypic Changes in Old *Cn* Cells

The upregulation of genes that encode metabolic pathways linked with FLC tolerance in aged *Cn* cells prompted an investigation into whether these changes align with phenotypic characteristics. We first assessed if the enhanced expression of genes associated with respiration in aged cells corresponds to increased ATP production. Confirming our hypothesis, old *Cn* cells exhibited elevated cellular ATP levels compared to their younger counterparts (Fig 2A). This observation was corroborated by heightened extracellular reduction of XTT (2,3-Bis(2-Methoxy-4-Nitro-5-Sulfophenyl)-5-[(Phenyl-Amino)Carbonyl]-2H-Tetrazolium Hydroxide), a result of NADH produced in the mitochondria (Fig S3). Quantification of mitochondrial mass (MM) using Mitotracker Green FM fluorescence also revealed a higher MM in old *Cn* cells (Fig 2B), indicating heightened mitochondrial respiration in old *Cn* cells.

**Figure 2.**
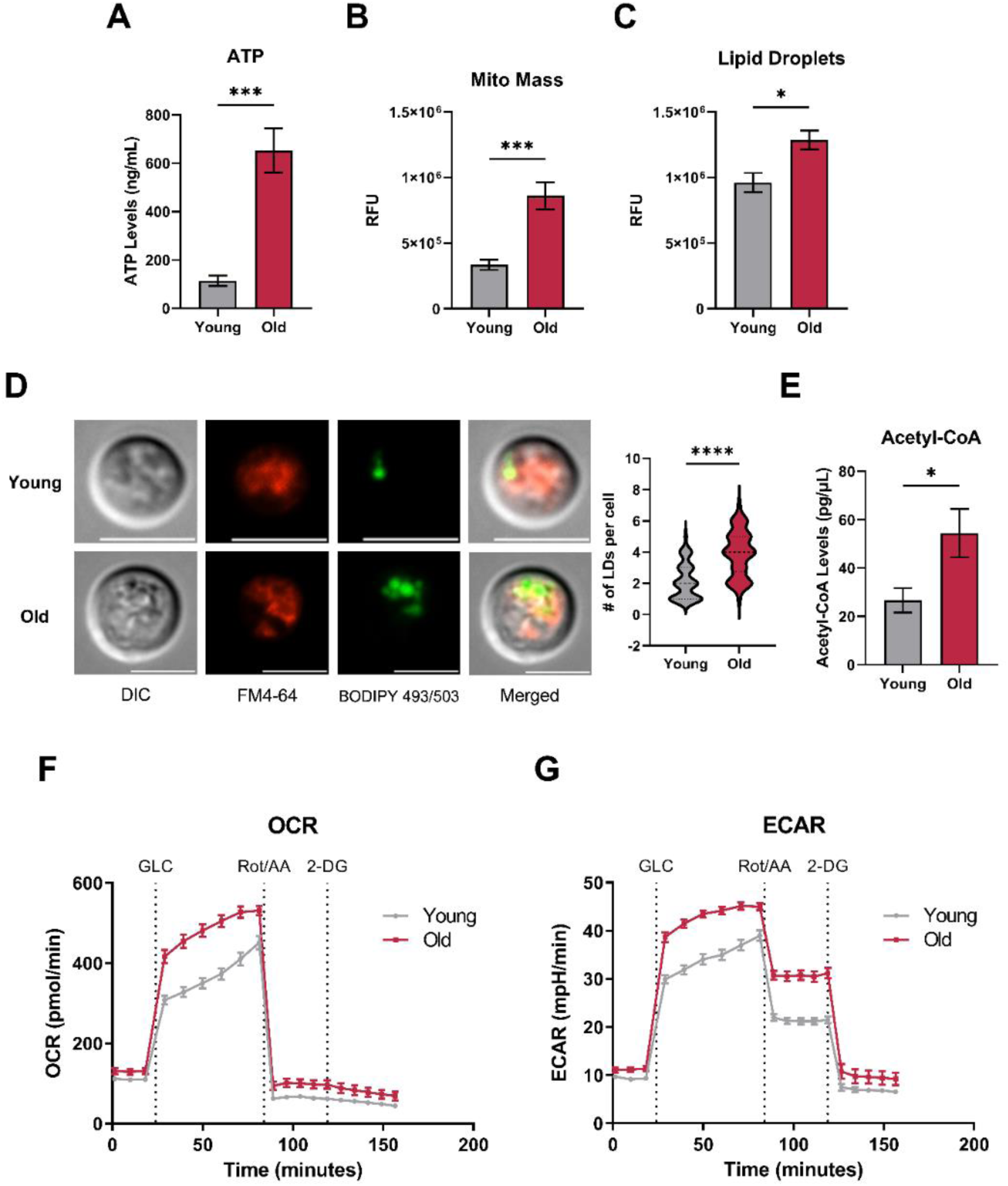
Old *Cn* cells Demonstrate Increased Energy Production and Metabolic Flux. (A) ATP levels of young and old cells were quantified using ATP Bioluminescent Assay. (B) Mitochondrial mass and (C) lipid droplets (LDs) of young and old cells were measured using a fluorescence plate reader by quantifying the relative fluorescent units (RFU) of Mitotracker Green FM and BODIPY 493/503 respectively. Bars signify the mean ± SEM of biological triplicates, and Student’s t-test was performed to determine the p-value (* p < 0.05, *** p < 0.001). (D) Vacuolar structure and LDs of young and old *Cn* cells were visualized by fluorescence microscopy after staining with FM 4-64 and BODIPY 493/503, respectively. Scale bars in white = 5µm. The number of LDs per cell (n = 50) was also counted, and Student’s t-test was performed to determine the p-value (**** p < 0.0001) (E) Acetyl CoA levels of young and old cells were quantified using Acetyl-Coenzyme A Fluorescent Assay. Bars signify the mean ± SEM of biological triplicates, and Student’s t-test was performed to determine the p-value (* p < 0.05). (F) Extracellular acidification rate (ECAR) and (G) oxygen consumption rate (OCR) profiles of young and old cells were generated from the Seahorse XF glycolysis stress test following injection of glucose (GLC), Rotenone/Antimycin A (Rot/AA), and 2-deoxyglucose (2-DG). Plotted values signify the mean ± SEM of 15 replicates.

Next, we explored the role of neutral lipids as potential energy sources via β-oxidation through the TCA cycle within the mitochondria. These lipids are stored in lipid droplets (LDs), which can be mobilized to the vacuole for energy. Quantifying LDs using BODIPY 494/503 fluorescence demonstrated an increase in old *Cn* cells compared to their younger counterparts (Fig 2C). Furthermore, microscopy revealed the localization of LDs to the vacuole, with an increased number of LDs per cell in aged cells (Fig 2D). This mobilization of LDs to the vacuole also supported the augmentation of acetyl-CoA pools through β-oxidation, as evidenced by the significant increase in acetyl-CoA levels in old *Cn* cells (Fig 2E).

To gain insights into the specific metabolic pathways contributing to the observed differences in energy production, we compared the metabolic flux between young and old *Cn* cells using the Seahorse XF Analyzer (Fig 2F and 2G). Basal extracellular acidification rate (ECAR) and oxygen consumption rate (OCR) were similar, indicating comparable levels of glycolysis, TCA cycle, and ETC activity under glucose-depleted conditions. However, following glucose injection, glycolysis and respiration increased more efficiently in old than in young *Cn* cells, as indicated by a higher rise in ECAR and OCR. When challenged with inhibitors of complex I and III (Rot/AA), old cells maintained higher glycolytic and TCA cycle activity in the absence of the ETC. Furthermore, adding glycolysis inhibitor 2-deoxyglucose (2-DG) led to a similar moderate decrease in OCR in both groups, indicating a reduction to baseline levels.

### Mitochondrial Morphology Dynamics in Old *Cn* Cells

The dynamic reorganization of the mitochondrial network, characterized by fission and fusion events, plays a pivotal role in responding to cellular energy demands (28, 29). We therefore investigated the impact of increased energy production in aged *Cn* cells on their mitochondrial morphology. Utilizing the mitochondrion-specific fluorescent dye, MitoTracker Green FM, and deconvolution fluorescence microscopy, we observed distinct differences in mitochondrial morphologies between young and old *Cn* cells, where young cells displayed diffuse mitochondrial networks while old cells exhibited highly fragmented mitochondria (Fig 3A).

**Figure 3.**
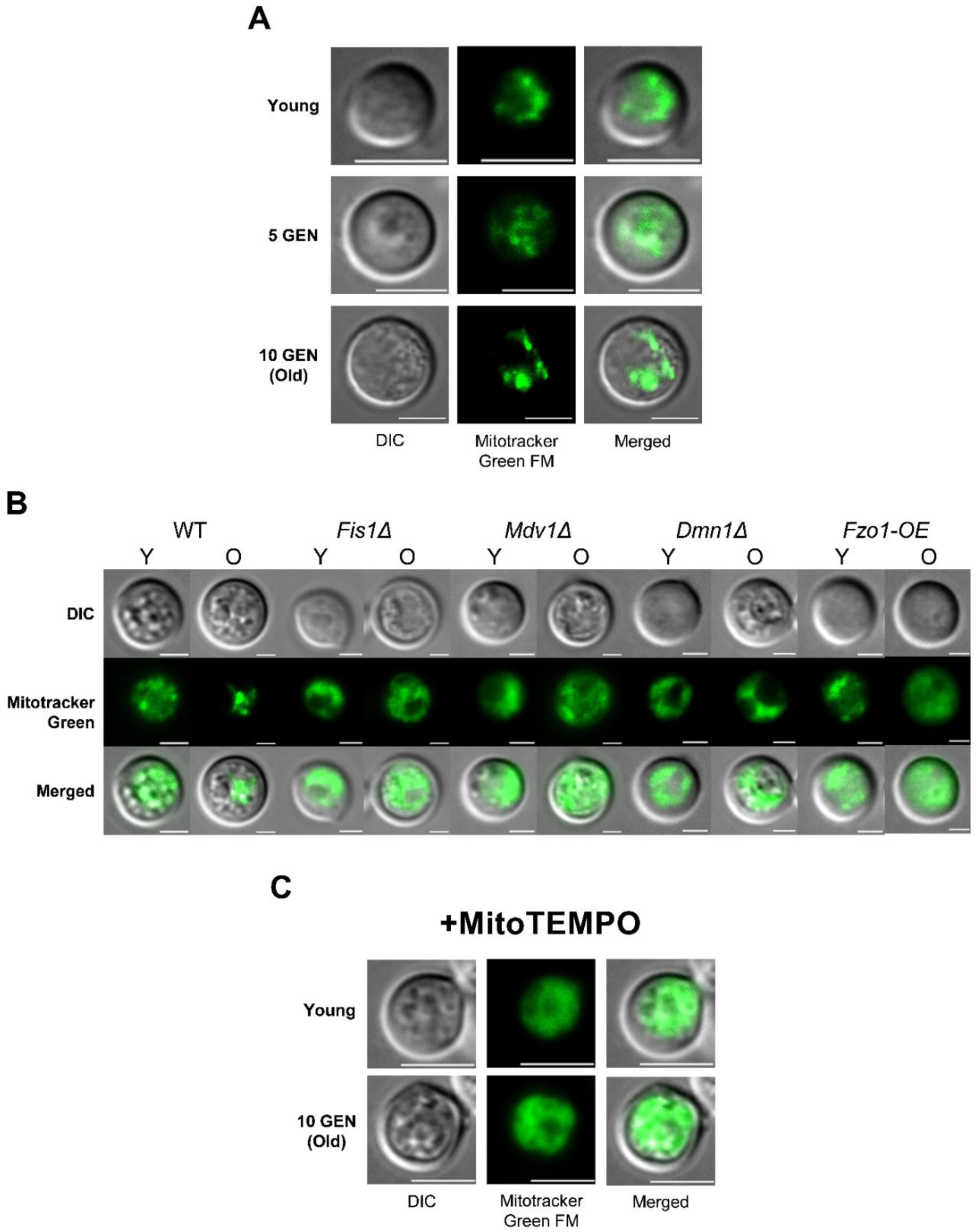
Dynamin-related proteins (DRPs) and Mitochondrial Reactive Oxygen Species Govern Age-associated Mitochondrial Fragmentation in Old *Cn* cells. **(A)** Mitochondrial morphologies of young (1-3 GEN), 5 GEN, and old (10 GEN) cells were visualized by fluorescence microscopy following Mitotracker Green FM staining. (B) Mitochondrial morphologies of young (Y) and old (O) cells of single-deletion strains for DRP fission genes (Δ*Fis1*, Δ*Mdv1*, and Δ*Dnm1*) and overexpression strain of the DRP fusion gene (*Fzo1-OE)* were also assessed using the same method. (C) Mitochondrial morphologies of young and old *Cn* cells grown in the presence of 50µM MitoTEMPO, a mitochondrially targeted antioxidant, were visualized using the same method. Scale bars in white = 5µm.

Given the distinct mitochondrial morphology in aged *Cn* cells, we explored the role of dynamin-related proteins (DRPs) in mediating mitochondrial fusion and fission(30). Genetic manipulations, including single-deletion strains for each of the fission genes (Δ*Fis1*, Δ*Mdv1*, and Δ*Dnm1*) and overexpression of the fusion gene (Fzo1-OE), revealed that these alterations influenced mitochondrial fragmentation in old cells (Fig 3B). Notably, defects in fission and increased fusion led to mitochondrial fragmentation, emphasizing the role of DRPs in age-associated changes.

Considering the observed increase in metabolic activity in old *Cn* cells, we hypothesized that the mitochondrial burden might contribute to the observed fragmentation. To test this hypothesis, we investigated whether mitochondrial oxidative stress, a byproduct of increased ATP production (31), played a role in mitochondrial fragmentation. Treatment with MitoTEMPO, a mitochondrially targeted antioxidant, resulted in similar mitochondrial morphologies in both young and old cells (Fig 3C), confirming that mitochondrial superoxide facilitates age-associated mitochondrial fragmentation in old cells.

### Age-associated Changes in ROS Levels, SOD Response, and Antioxidant Defense Mechanisms

The increased mitochondrial burden resulting from ATP production in aged *Cn* cells prompted a deeper exploration of the mitochondrial stress response. First, we assessed cellular reactive oxygen species (cROS) and mitochondrial reactive oxygen species (mROS) levels, indicative of oxidative stress resulting from ATP production through oxidative phosphorylation. Old *Cn* cells exhibited elevated H_2_DCFDA and MitoSox Red fluorescence, signifying increased cROS (Fig 4A) and mROS (Fig 4B), respectively. Furthermore, a higher mitochondrial calcium level in old *Cn* cells, as indicated by increased Fluo-4 AM fluorescence (Fig 4C).

**Figure 4.**
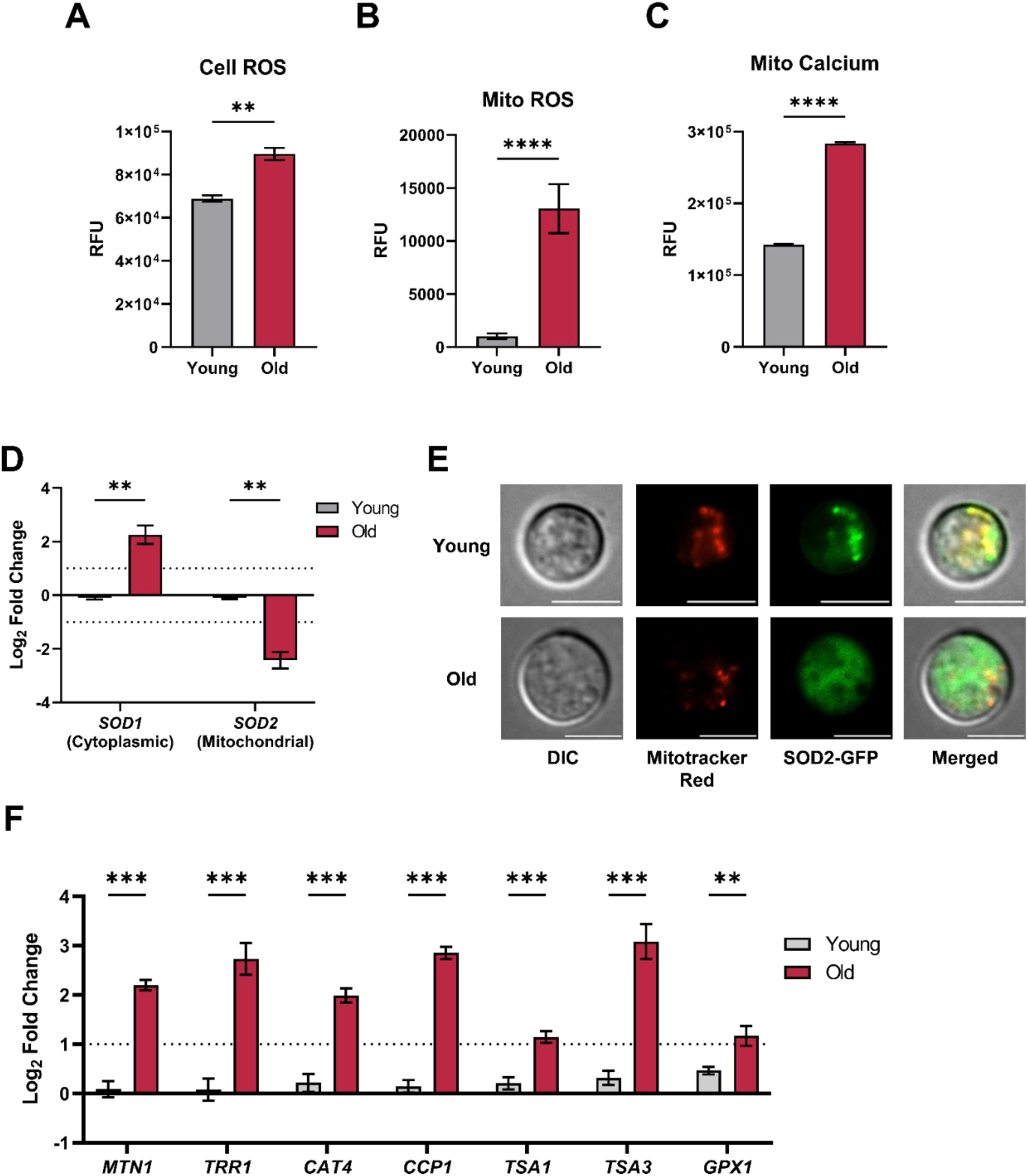
A Suite of Cellular Antioxidant Defense Mechanism Regulates Oxidative Stress in Old *Cn* Cells. (A) Cellular reactive oxygen species (ROS), (B) mitochondrial ROS, and (C) mitochondrial calcium levels of young and old cells were measured using a fluorescence plate reader by quantifying the fluorescent intensity of H_2_DCFDA, MitoSox Red, and Fluo-4 AM respectively. Bars signify the mean ± SEM of biological triplicates, and Student’s t-test was performed to determine the p-value (** p < 0.01, *** p < 0.001, **** p < 0.0001). (D) mRNA expression of superoxide dismutase genes (*SOD1* and *SOD2*) was analyzed by RT-qPCR in young (grey bars) and old (red bars) cells. Bars signify the mean ± SEM of biological triplicates and Student’s t-test was performed to determine the p-value (** p < 0.01). The mRNA expression is presented as Log_2_FC relative to the expression of *ACT1*, which was used as an internal control. (E) Cellular localization of SOD2-GFP in young and old cells was visualized by fluorescence microscopy. Cells were stained with Mitotracker Red to check the co-localization of SOD2-GFP with the mitochondria. Scale bars in white = 5µm. (F) mRNA expression of antioxidant genes was analyzed by RT-qPCR in young (grey bars) and old (red bars) cells. Bars signify the mean ± SEM of biological triplicates, and Student’s t-test was performed to determine the p-value (** p < 0.01, *** p < 0.001). The mRNA expression is presented as Log_2_FC relative to the expression of *ACT1*, which was used as an internal control.

A crucial mechanism in response to elevated ROS is the action of superoxide dismutase (SOD), which converts ROS to harmless water and free oxygen (32). In *Cn*, Sod1 and Sod2 regulate cROS and mROS, respectively (33, 34). RT-qPCR analysis revealed the upregulation of *SOD1* and downregulation of *SOD2* in old *Cn* cells (Fig 4D). Intriguingly, the intracellular localization of Sod2-GFP in old cells showed dispersed cytoplasmic distribution, contrasting with the predominantly mitochondrial localization in young *Cn* cells (Fig 4E).

To reconcile the downregulation of *SOD2* as well as cytoplasmic localization of Sod2-GFP despite the high mROS in old *Cn* cells, we explored the regulation of other components of the cellular antioxidant defense system. Our findings revealed an upregulation of genes encoding copper-detoxifying metallothionein 1 (*MTN1*), thioredoxin reductase (*TRR1*), catalase 4 (*CAT4*), mitochondrial cytochrome c peroxidase (*CCP1*), thiol peroxidase 1 and 3 (*TSA1*, *TSA3*), and glutathione peroxidase 1 (*GPX1*) in old *Cn* cells (Fig 4F). Expression levels of other genes encoding other components of the cellular antioxidant defense system were unchanged (Fig S4).

### Role of ABC Transporters and Ergosterol Dynamics in Age-associated FLC Tolerance

The development of FLC resistance in *Cn* has been linked to enhanced expression and activity of efflux pumps, resulting in decreased intracellular drug concentrations (14, 15). Given their involvement in drug efflux, we investigated the potential contribution of ABC transporters to age-associated FLC tolerance in old *Cn* cells. Quantification of the mRNA expression of key ABC transporters revealed a significant upregulation of *AFR1*, *AFR2*, *AFR3*, and *MDR1* in old cells (Fig 5A). To further confirm the potential role of the ABC transporters in age-associated FLC tolerance, we assessed their efflux activity using fluorescent probes Nile red (NR) and Rhodamine 6G (R6G) (35). The results demonstrated heightened efflux of both NR and R6G in old *Cn* cells (Fig 5B and 5C), indicating increased activity of ABC transporters in aged cells.

**Figure 5.**
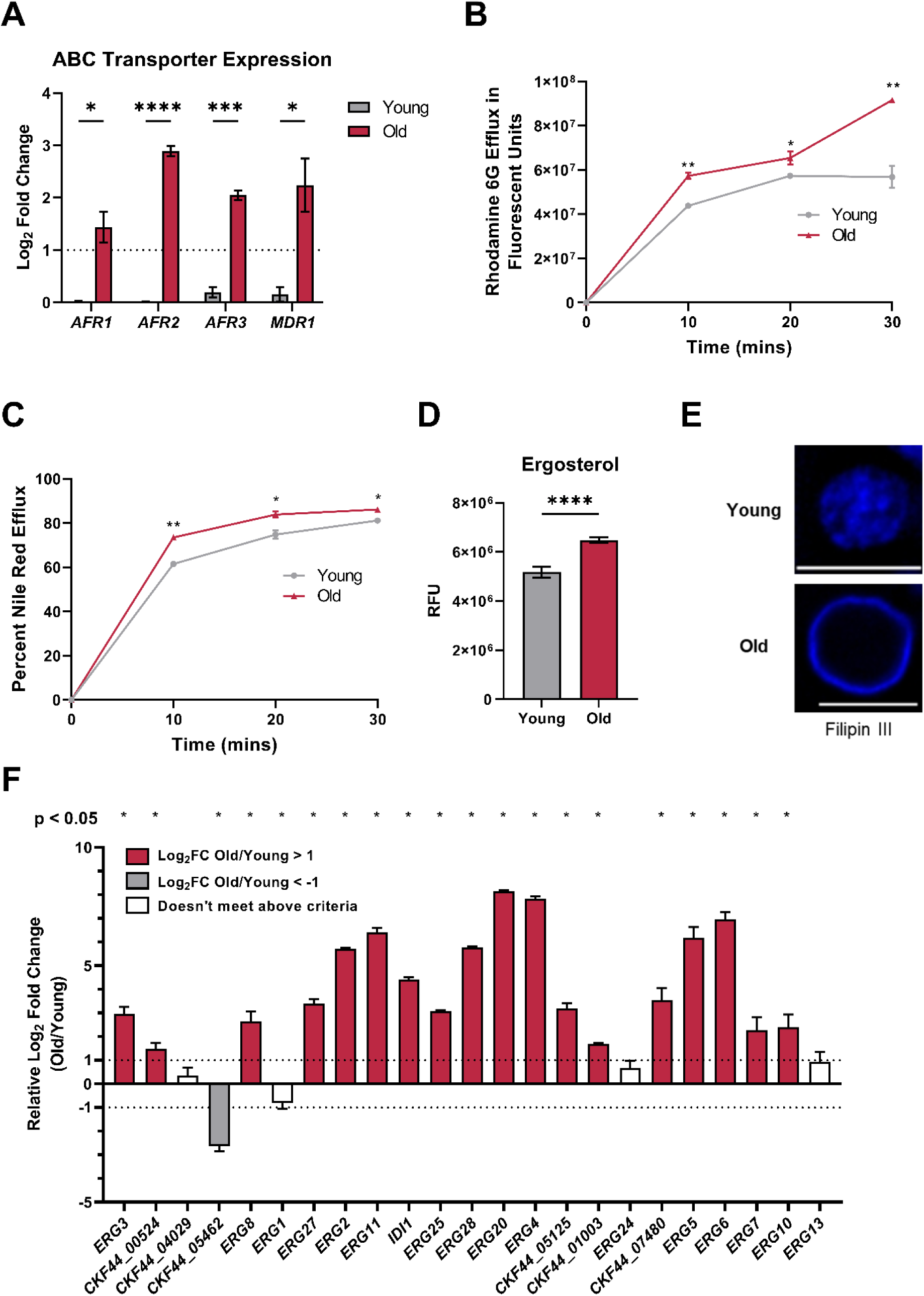
Old *Cn* Cells Display Changes to ATP-Binding Cassette (ABC) Transporters and Ergosterol Synthesis. (A) mRNA expression of genes encoding ABC transporters was analyzed by RT-qPCR in young (grey bars) and old (red bars) cells. The mRNA expression is presented as Log2FC relative to the expression of ACT1, which was used as an internal control. Bars signify the mean ± SEM of biological triplicates, and Student’s t-test was performed to determine the p-value (* p < 0.05, *** p < 0.001, **** p < 0.0001). ABC transporter activities of young and old cells were measured by quantifying the efflux of (B) Rhodamine 6G and (C) Nile Red using a fluorescence plate reader over 30 minutes. Markers signify the mean ± SEM of biological triplicates (* p < 0.05, ** p < 0.01). (D) Sterol levels of young and old cells were measured using a fluorescence plate reader by quantifying the fluorescent intensity of Filipin III. Bars signify the mean ± SEM of biological triplicates. (**** p < 0.0001) (E) Sterol structure of young and old cells was visualized by fluorescence microscopy following staining with Filipin III. Scale bars in white = 5µm. (F) mRNA expression of ergosterol synthesis genes was analyzed by RT-qPCR in young and old cells, and mean Log2FC comparing expression in old/young cells (with error bars reflecting ± SEM) is shown for each gene. Genes in red indicate upregulation in old with Log2FC > 1, while genes in grey indicate downregulation in old with Log2FC < −1. Bars signify the mean ± SEM of biological triplicates. Student’s t-test was performed to determine the p-value for the differences in mRNA expression in young and old cells for each gene, as marked above the graph.

Considering that changes in ergosterol levels also contribute to FLC resistance, we explored ergosterol dynamics in young and old *Cn* cells. Using filipin, a fluorescent dye that binds to membrane sterols (36), we observed elevated fluorescence in old cells, suggesting higher cellular ergosterol levels (Fig 5D). Additionally, fluorescence microscopy revealed distinct intracellular staining patterns, with old cells displaying a ring-like pattern in contrast to the diffuse pattern in young cells (Fig 5E). To understand the molecular basis of altered ergosterol levels, we examined the mRNA expression of genes involved in the ergosterol synthesis pathway. Out of 23 genes, old cells exhibited significant upregulation of 18 genes (Log_2_FC Old/Young ≥ 1) and downregulation of 1 gene (Log_2_FC Old/Young ≤ −1) (Fig 5F), emphasizing the regulatory changes in ergosterol synthesis during aging.

### Mitochondrial Signaling Dynamics Contributing to Age-associated FLC Tolerance

Mitochondria, beyond their canonical role in ATP production, orchestrate cellular responses, engaging in intricate crosstalk with various organelles and cellular pathways (18). Considering the distinctive mitochondrial characteristics exhibited by aged *Cn* cells, we tested the possibility that the mitochondrial stress exhibited by old cells leads to downstream cellular signaling. Drawing insights from previous studies demonstrating the influence of mitochondrial signaling on drug resistance in fungal species (16, 17), we hypothesized that the observed mitochondrial stress in aged *Cn* cells might trigger downstream signaling pathways contributing to their FLC tolerance.

To test this hypothesis, we first probed the impact of antioxidants on FLC tolerance by examining the specific effects of ascorbic acid (AA) and glutathione (GSH). Both young and old cells treated with antioxidants exhibited diminished cROS levels compared to untreated controls (Fig S5A). While these antioxidants significantly reduced FLC killing in young cells, they did not affect FLC killing in aged cells (Fig 6A). Further exploration involved targeted scavenging of mitochondrial superoxide using MitoTEMPO. While MitoTEMPO generally attenuated FLC killing of young cells (Fig 6B), the response in aged cells proved to be more complex. Contrary to our expectations, aged cells exhibited increased FLC killing in the presence of MitoTEMPO, surpassing the effect observed in young cells. We also confirmed that young and old cells treated with MitoTEMPO displayed diminished mROS levels compared to untreated controls (Fig S5B).

**Figure 6.**
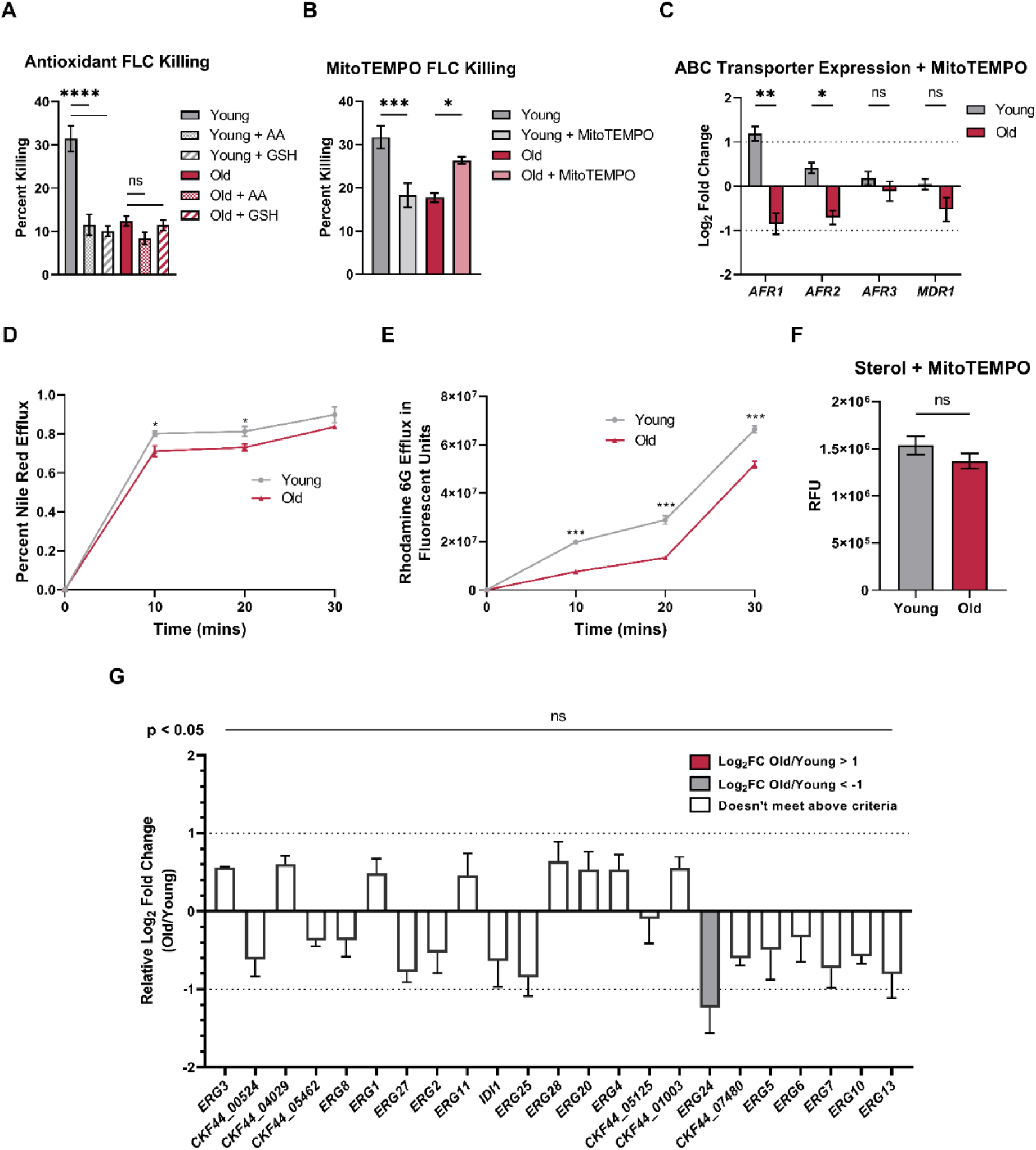
mROS Contributes to Age-associated Fluconazole Tolerance in Old *Cn* Cells. Resilience to fluconazole (FLC) killing was measured in young and old cells (A) ± 10mM ascorbic acid (AA) and 10mM glutathione (GSH) and (B) ± 50µM MitoTEMPO by calculating the change in colony-forming units (CFU) after treatment with 50μg/mL FLC for 3 hours. Bars signify the mean ± SEM of biological triplicates. One-way ANOVA with Tukey’s multiple comparisons test was used to determine the p-value (ns p > 0.05, * p < 0.05, *** p < 0.001, **** p < 0.0001). (C) mRNA expression of genes encoding ABC transporters was analyzed by RT-qPCR in young (grey bars) and old (red bars) cells grown in the presence of 50 µM MitoTEMPO. The mRNA expression is presented as Log2FC relative to the expression of ACT1, which was used as an internal control. Bars signify the mean ± SEM of biological triplicates, and Student’s t-test was performed to determine the p-value (ns p > 0.05, * p < 0.05, ** p < 0.01). ABC transporter activities of young and old cells grown in the presence of 50 µM MitoTEMPO were measured by quantifying the efflux of (D) Rhodamine 6G and (E) Nile Red using a fluorescence plate reader over 30 minutes. Markers signify the mean ± SEM of biological triplicates and Student’s t-test was performed to determine the p-value (* p < 0.05, *** p < 0.001). (F) Sterol levels of young and old cells grown in the presence of 50µM MitoTEMPO were measured using a fluorescence plate reader by quantifying the fluorescent intensity of Filipin III. Bars signify the mean ± SEM of biological triplicates and Student’s t-test was performed to determine the p-value (ns p > 0.05). (G) mRNA expression of ergosterol synthesis genes was analyzed by RT-qPCR in young and old cells, and mean Log2FC comparing expression in old/young cells grown in the presence of 50µM MitoTEMPO (with error bars reflecting ± SEM) is shown for each gene. Genes in red indicate upregulation in old with Log2FC > 1, while genes in grey indicate downregulation in old with Log2FC < −1. Bars signify the mean ± SEM of biological triplicates. Student’s t-test was performed to determine the p-value for the differences in mRNA expression in young and old cells for each gene, as marked above the graph.

These results prompted a deeper inquiry into how mitochondrial superoxide influences age-associated FLC tolerance. First, we tested the possibility that MitoTEMPO treatment alters the metabolic upregulation that we observed in old cells. However, Seahorse Analysis revealed that MitoTEMPO does not affect the metabolic flux that was previously observed in young and old cells (Fig S6A and S6B). Next, we probed how the addition of MitoTEMPO influences the changes in ABC transporter expression and increased efflux seen in old cells. When the cells underwent replicative aging in the presence of MitoTEMPO, old cells showed a slight downregulation of *AFR1* and *AFR2* compared to young cells, while there were no significant differences in *AFR3* and *MDR1* expression (Fig 6C). These transcriptional changes were consistent with efflux activity, as young cells demonstrated higher efflux activity measured by fluorescent probes NR and R6G (Fig 6D and 6E).

We also examined whether mROS influences age-associated changes in ergosterol regulation. We observed no significant difference in fluorescence in young and old cells grown in the presence of MitoTEMPO (Fig 6F), suggesting similar cellular ergosterol levels. Upon further interrogation, we found no significant differential expression of genes involved in the ergosterol synthesis pathway between young and old cells grown in the presence of MitoTEMPO (Fig 6G). Taken together, the loss in age-dependent changes in ABC transporters and ergosterol synthesis emphasize the importance of mROS in mediating these changes to drive age-associated FLC tolerance in *Cn*.

## DISCUSSION

This is the first study to interrogate the mechanisms by which replicative aging confers increased FLC tolerance in *Cn*. Ten-generation-old *Cn* cells demonstrated widespread transcriptional changes that encompass various aspects of metabolism. These cells displayed increased mitochondrial activity that resulted in increased ATP production, which fuels ABC transporters. These changes are accompanied by the upregulation of ergosterol synthesis and ABC transporters, which are dependent on the mROS signaling resulting from increased mitochondrial stress. These findings constitute the mechanistic basis that underlies the age-associated FLC tolerance in old *Cn* cells (Fig 7).

**Figure 7.**
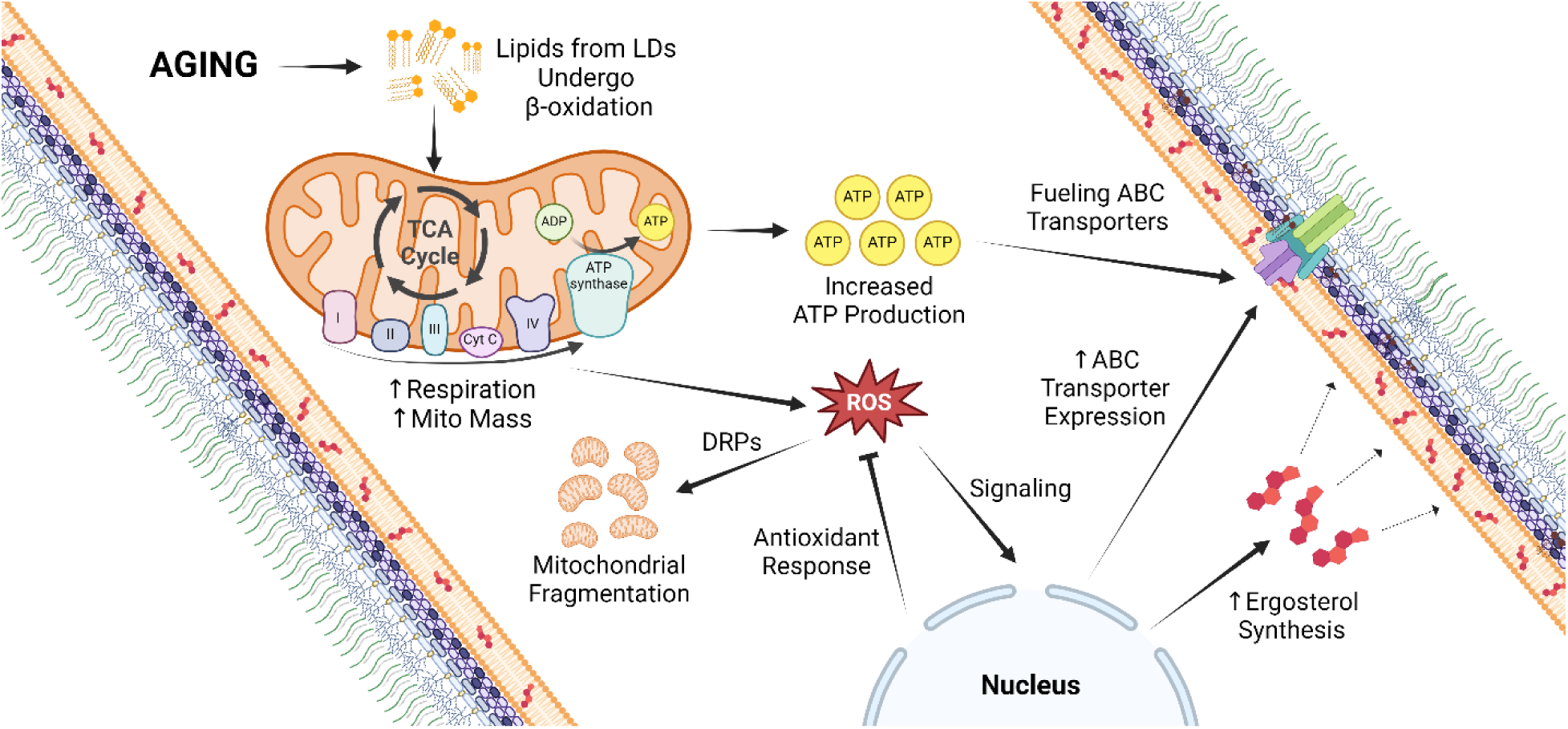
Metabolic Changes in the Mitochondria Drive Age-associated FLC Tolerance in *Cn*. Replicative aging leads to increased use of lipids from lipid droplets (LDs) as potential energy sources by β-oxidation through the tricarboxylic acid (TCA) cycle within the mitochondria. The heightened respiration and higher mitochondrial mass result in increased ATP production. The increased mitochondrial activity leads to the production of reactive oxygen species (ROS), an unavoidable byproduct of respiratory chain function during ATP synthesis. Increased ROS production leads to mitochondrial fragmentation, which is mediated by dynamin-related proteins (DRPs), and activation of a suite of antioxidant responses. Increased mitochondrial ROS also drives FLC tolerance by increasing ergosterol synthesis and upregulating ATP-binding cassette (ABC) transporters. Increased ATP production fuels ABC transporters, which further contributes to FLC tolerance. Created with BioRender.com.

Using GO and KEGG enrichment analyses, we were able to classify differentially expressed genes according to their molecular function, biological processes, and biological pathways. Genes encoding proteins involved in transition metal ion transport were upregulated in old *Cn* cells. Transition metals such as copper, iron, and zinc play a central role in fungal pathogenesis (37). Additionally, genes involved in β-glucan and polysaccharide metabolic processes were also upregulated in old *Cn* cells. β-glucan is the most important structural component of the fungal cell wall and represents 50–60% of the dry weight of this structure (38). *Cn* mutants lacking genes involved in β-1,6-glucan synthesis are more sensitive to environmental and chemical stress and display important alterations in the cell wall composition leading to the loss of virulence in the mammalian host (39). Lastly, the polysaccharide capsule serves as the outer protective layer against various stressors and is one of the major cryptococcal virulence factors (40). These results highlight the overlap of the functional groupings between the upregulated genes in old *Cn* cells from this study and genes that facilitate cellular processes that confer enhanced fitness.

To complement the results from our transcriptomic analyses, we looked to identify phenotypic characteristics that were consistent with the upregulation of cellular respiration through fatty acid oxidation. The ability of *Cn* cells to produce energy through respiration is crucial for cryptococcal virulence*, in-vivo* resilience, and antifungal resistance (41, 42). We found that older *Cn* cells produce higher intracellular ATP levels. This result was also confirmed by the XTT assay, a viability staining method that can serve as a marker for mitochondrial activity based on the reduction of XTT by mitochondrial dehydrogenase. Lastly, older *Cn* cells demonstrated higher mitochondrial mass compared to younger cells, further confirming characteristics consistent with their high ATP levels.

The majority of ATP is synthesized in the mitochondria during respiration and fatty acid oxidation. Neutral lipids are stored in LDs, and hydrolysis of LDs provides an extra source of energy through fatty acid oxidation (43). LDs communicate via direct contact with other cellular organelles, and our data demonstrates that *Cn* LDs localize to the vacuole regardless of the cells’ replicative age. In yeast, the vacuole functions as the major catabolic organelle and interacts with LDs to regulate lipid homeostasis (43). Notably, older cells exhibited a higher amount of total intracellular LDs as well as a larger number of LDs per cell compared to young cells. Furthermore, we observed an increase in intracellular acetyl-CoA levels in the old cells, indicating higher fatty acid oxidation. Altogether, these results indicate that older *Cn* cells have a higher cellular supply of energy sources that can respond to cellular metabolic demands.

The Seahorse Analysis also revealed some notable changes in metabolic flux in old *Cn* cells. First, the old cells demonstrated a higher maximal level of glycolysis and respiration, as well as a higher rate of response to glucose. These are supported by the higher OCR/ECAR maximum in old cells and the higher increase in OCR/ECAR levels immediately following glucose injection. These differences indicate that older cells sense and transport glucose faster than young cells. Additionally, older cells may have a higher metabolic turnover and demand, leading them to respond to nutrients faster and at a higher maximum capacity than young cells. Another notable difference was that old cells also showed significantly higher glycolysis following the inhibition of mitochondrial oxidative phosphorylation by Rot/AA. Because residual glycolysis at this stage is dependent on the NAD+ levels in *Cn* (44), it could be that a larger NAD+ pool in old *Cn* cells drives glycolysis to meet the increased energy demand.

To maintain proper functionality, the mitochondrial network undergoes constant dynamic reorganization that leads to changes in mitochondrial morphologies (30). During healthy growth, mitochondria exhibit a diffuse morphology. However, under various cellular stresses, mitochondria can undergo fusion and fission, leading to tubular and fragmented mitochondrial morphologies, respectively (45). These morphological changes are facilitated by DRPs, where fission proteins Fis1, Mdv1, and Dmn1 mediate mitochondrial fragmentation (30). We observed progressive fragmentation of the mitochondria upon replicative aging, where young cells displayed diffuse mitochondrial networks while old cells exhibited highly fragmented mitochondria. In contrast, we did not observe age-associated mitochondrial fragmentation in aged in fission-defective strains *(ΔDnm1, ΔMdv1,* and *ΔFis1)*, indicating that all three fission proteins are necessary to mediate mitochondrial fragmentation seen in aging.

Studies have shown that an increase in oxidative stress during replicative aging results in mitochondrial network fragmentation (46, 47). This increase in oxidative stress originates from ROS, an unavoidable byproduct of respiratory chain function during ATP synthesis in a eukaryotic cell. Indeed, we observed higher cROS as well as higher mROS in older *Cn* cells than in younger cells. Quenching the mROS by adding MitoTEMPO during the aging process resulted in similar mitochondrial morphologies between young and old cells, indicating that the increased mROS production during replicative aging contributes to the mitochondrial fragmentation observed in old *Cn* cells. This was further supported by the high mitochondrial calcium levels, as mitochondrial calcium overload can also be a form of mitochondrial stress that induces mitochondrial fragmentation (48).

SOD serves as an important antioxidant defense by catalyzing the breakdown of ROS to minimize the cellular damage caused by the generation of superoxide during respiration (49). While we observed upregulation of *SOD1* in response to higher cROS in old *Cn* cells, we also detected downregulation of *SOD2* despite the elevated mROS. Furthermore, we saw cytoplasmic localization of Sod2 in older cells, contrasted with mitochondrial localization in young cells. We also detected a widespread upregulation of other antioxidant response proteins in old cells. Some of these proteins, such as Mtn1, Trr1, Ccp1, and Tsa3 are known to localize to the mitochondria (50–52). The presence of this redundancy in the mitochondrial antioxidant defense system might explain the downregulation of *SOD2* and cytoplasmic localization of Sod2 seen in old *Cn* cells. Notably, we also observed an upregulation of Cat4 and Gpx1, which both catalyzes the further detoxification of hydrogen peroxide, produced by SOD from superoxide, into water and oxygen.

The efflux pumps that are associated with FLC resistance in *Cn* belong to the ABC transporter family, which requires an adequate supply of ATP for FLC efflux (14, 15, 53–55). Afr1 is the dominant membrane transporter in *Cn*, while Afr2, Afr3, and Mdr1 have also been shown to play a role in azole resistance (14, 15, 55). Consistent with our previous study demonstrating that older *Cn* cells are more resistant to FLC (15), we saw significant upregulation of all four ABC transporters in these cells. Using fluorescent probes NR and R6G to measure efflux activity, we validated the upregulation of ABC transporters in older cells. This difference in NR and R6G efflux activities is also consistent with the higher ATP levels seen in old cells, which serve as fuel to drive these cellular efflux pumps.

Disruptions to the ergosterol metabolic pathway have been implicated in both diminished effectiveness of azole compounds (12, 13). Using Filipin III, a sterol-binding fluorescent dye, we observed an increased ergosterol content in old *Cn* cells. Consistent with these findings, we also observed significant upregulation of 18 out of 23 genes involved in the ergosterol synthesis pathway. Notably, *ERG11*, encoding the enzyme that is directly targeted by FLC (7) and *ERG6*, which is crucial for *Cn* virulence by affecting membrane integrity and dynamics (56), were upregulated. Ergosterol biosynthesis starts with the condensation of two acetyl-CoA molecules (57), and ergosterol can be stored in LDs in the form of steryl ester (58). Interestingly, we also found higher levels of acetyl-CoA and LDs in old cells, further supporting our results that old cells upregulate ergosterol synthesis.

To examine the potential role of increased ROS in mediating the age-associated FLC tolerance in old *Cn* cells, we tested the FLC tolerance of old cells that have undergone replicative aging in the presence of various antioxidants. General antioxidants like AA and GSH significantly reduced FLC susceptibility in young cells but had no effect in aged cells. This observation is consistent with previous studies that have shown that ROS contributes to FLC-mediated growth inhibition, which can be rescued by using these antioxidants (59–61). Interestingly, aged *Cn* cells in the presence of MitoTEMPO exhibited increased susceptibility to FLC, suggesting that oxidative stress response, which mediates age-associated FLC resistance in old cells might be specific to mROS.

Old cells that have undergone replicative aging in the presence of MitoTEMPO, compared to their younger counterparts, displayed downregulation of *AFR1* and *AFR2* and no significant differential regulation of *AFR3* and *MDR1*. These transcriptional changes were also supported by NR and R6G studies demonstrating the decreased efflux activity of these cells. Furthermore, these MitoTEMPO-treated old *Cn* cells exhibited no difference in ergosterol content and no longer displayed widespread significant upregulation of ergosterol synthesis genes. These distinct changes in ABC transporter and ergosterol phenotypes brought on by scavenging superoxide confirmed that mROS plays a pivotal role in mediating the observed age-associated characteristics that contribute to the resistance of old *Cn* cells against FLC.

Several studies have highlighted the role of mitochondria-driven signaling in mediating FLC resistance in pathogenic fungi. In *Candida glabrata*, the loss of mitochondrial function is associated with the acquisition of azole resistance through the upregulation of genes encoding ABC transporters (17, 62). In *Saccharomyces cerevisiae* and *Aspergillus fumigatus*, mitochondrial dysfunction arising from alteration of cofilin residues has been shown to trigger retrograde signaling, resulting in the upregulation of ABC transporter genes and activation of fatty acid β-oxidation to increase acetyl-CoA and ATP levels (63, 64). In *Cn*, Mar1 has been shown to mediate FLC tolerance by altering cellular mitochondrial metabolism (65). Additionally, transcriptomic analysis of *Cn* during exposure to oxidative stress revealed that there is an induction of an antifungal drug resistance response upon the treatment of *Cn* with hydrogen peroxide (66). These studies support our conclusions that increased mROS promotes FLC tolerance in old *Cn* cells.

This is the first study that highlights the mechanism that underlies age-associated FLC tolerance in *Cn* cells of advanced replicative age. Replicative aging results in a heterogeneous cell population composed of cells with varying generational ages. Cells of advanced generational age accumulate during *in-vivo Cn* and *C. glabrata* infection, indicating that they are selected by the host environment (25, 26). We demonstrate that aging in *Cn* cells modulates mitochondrial metabolism and promotes FLC tolerance, ultimately contributing to their resilience. Genetic antifungal resistance is uncommon among *Cn* (10, 11), and do not explain the poor clinical outcomes that are observed with cryptococcal meningitis (67). Our findings may shed light on why standard antifungal resistance testing performed with *in-vitro*-grown *Cn* populations does not predict treatment failure. Understanding the non-genetic age-dependent heterogeneity in fungal populations not only expands our comprehension of drug tolerance but also greatly advances our endeavors in identifying novel therapeutic targets.

## METHODS

### Strains and media

*Cn* strains were kept on YPD agar plates. Synthetic media (68) was used for the cultivation of yeast cells. 10mM ascorbic acid, 10mM glutathione, and 50μM MitoTEMPO were added to the media when needed. All the strains used in this study are listed in Table S1.

### Isolation of Young and Old *Cn* Cells

Isolation of young (0-3 generations) and old (10 generations) *Cn* cells was performed as previously described (69).

### RNA-seq Analysis

RNA was extracted using the RNAeasy Plus kit, following the manufacturer’s guidelines. RNASeq was performed by Novogene Co. Messenger RNA was purified from total RNA using poly-T oligo-attached magnetic beads and converted to cDNA. Libraries were clustered and sequenced on an Illumina NovaSeq 6000 platform, which utilizes a paired-end 150bp sequencing strategy according to the manufacturer’s instructions. Raw data (raw reads) of fastq format were firstly processed through in-house perl scripts, where clean data (clean reads) were obtained by removing reads containing adapter, reads containing ploy-N and low quality reads from raw data. At the same time, Q20, Q30 and GC content the clean data were calculated. All the downstream analyses were based on the clean data with high quality. Differential expression analysis was performed using the DESeq2Rpackage (1.20.0). The resulting P-values were adjusted using the Benjamini and Hochberg’s approach for controlling the false discovery rate. Genes with an adjusted P-value ≤ 0.05 and |log_2_FoldChange| ≥ 1 were assigned as differentially expressed. Gene ontology (GO) Enrichment analysis was performed by running differentially expressed genes through FungiDB, selecting for Biological Process (BP) and Molecular Function (MF) with a p-value cutoff of 0.05. Kyoto Encyclopedia of Genes and Genomes (KEGG) Analysis was performed by using clusterProfiler R package to test the statistical enrichment of differential expression genes in KEGG pathways. Volcano plot and heatmaps were generated by Novogene Co.

### Measurement of Cellular ATP Levels

Cellular ATP levels were quantified using ATP Bioluminescent Assay Kit as previously described (68). The assay was performed in triplicate.

### XTT Assay

Cellular XTT [2,3-bis(2-methoxy-4-nitro-5-sulfophenyl)-2H-tetrazolium-5-carboxanilide] levels were quantified using XTT Assay Kit following the manufacturer’s guidelines. Briefly, 5×10^6^ young and old cells were washed three times with SM and resuspended in 500μL SM. For each reaction, 150μL activated XTT solution was added and was incubated at 37°C for 3 hours protected from light. Following incubation, cells were pelleted, and the supernatant fraction was measured for absorbance at 475nm with the spectrophotometer. Heat-killed cells (5 mins incubation at 65°C) were used as negative controls. The assay was performed in triplicate.

### Measurement of Cellular Acetyl-CoA Levels

10^7^ cells were mixed with 400μL assay buffer and vortexed for 15 min in the presence of sterile acid-washed glass beads to initiate lysis. Following lysis, samples were retrieved by removing the cellular debris and beads by centrifugation. 50μL samples were loaded in black 96-well plates and 10μL of Acetyl-CoA Quencher was added to each sample and incubated at room temperature for 5 minutes. Then, 2μL of Quench Remover was added and incubated for an additional 5 minutes. Samples were mixed with 2μL Acetyl-CoA Substrate Mix, 1μL Conversion Enzyme, 5μL Acetyl-CoA Enzyme Mix, 2μL Fluorescent Probe. Reaction was incubated for 10 minutes at 37°C, and fluorescence was read in a fluorescent plate reader with λex = 535nm/λem = 587nm. Acetyl-CoA standard solution was used in serial dilution to generate a standard curve, which was used to interpolate the cellular Acetyl-CoA levels. The assay was performed in triplicate.

### Quantification of Cellular Phenotypes Using Fluorescent Dyes

Young and old cells (10^6^) were stained protected from light with the following dyes with the respective concentrations and incubation times/temperature: 10µM H_2_DCFDA for 30 mins at 37°C, 500nM MitoSOX Red for 15 mins at 37°C, 200nM MitoTracker Green FM for 30 mins at 37°C, 5µM BODIPY 493/503 for 30 mins at 24°C, and 5μg/mL Filipin III for 5 mins at 24°C. After staining, the cells were washed three times, resuspended in PBS, and 200µL of washed stained cells were loaded in black 96-well plates. Fluorescence was read in a fluorescent plate reader with the following wavelengths: λex = 495nm/λem = 525nm for H_2_DCFDA, λex = 510 nm/λem = 580 nm for MitoSOX Red, λex = 490nm/λem = 516nm for MitoTracker Green FM, λex = 491nm/λem = 516nm for BODIPY 493/503, and λex = 360nm/λem =470nm for Filipin III. Fluorescence was measured in relative fluorescence intensity (RFI) by subtracting the RFI values between the stained and non-stained cells. Experiments were performed in biological triplicate.

### Fluorescence Microscopy

Fungal cells were stained with MitoTracker Green FM, BODIPY 493/503, Filipin III (staining conditions described above), 10μg/mL FM 4-64 for 10 mins at 37°C, and 500nM MitoTracker Red FM for 30 mins at 37°C. After staining, cells were fixed and imaged as previously described (68). Each experiment was repeated at least once on different days under the same conditions.

### Seahorse XF Analysis

A Glycolytic Rate assay was performed on a Seahorse Biosciences XFe96 extracellular flux analyzer for oxygen consumption rate (OCR) and extracellular acidification rate (ECAR) as previously described (44) with minor modifications. Briefly, cartridges were hydrated overnight in Agilent Seahorse XF calibrant. 180μL of young and old cells (2 x 10^6^ cells/mL) was loaded into each well. Assay solutions were injected to a final concentration of 20 mM glucose, 3.5μM Rotenone/Antimycin A (Rot/AA), and 100mM 2-deoxyglucose (2-DG). Cells and solutions were prepared on Seahorse XF Base Medium supplemented with 2mM L-glutamine and 5mM HEPES pH 7.4. Each condition was measured with at least 15 replicates.

### RT-qPCR Analysis

RNA extraction, cDNA conversion, and qPCR were performed following the manufacturers’ guidelines as previously described (15). The oligonucleotides used in this study are described in Table S2. The assay was performed in biological triplicate.

### Nile Red (NR) and Rhodamine 6G (R6G) Efflux Assays

NR and R6G assays were performed to analyze the efflux activities of the membrane transporters as previously described (15, 68). The assays were performed in biological triplicate.

### Fluconazole (FLC) Killing Assay

FLC killing assay was performed as previously described (15). Percent killing was analyzed by the following formula:

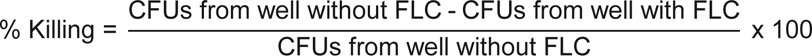

The assay was performed in biological triplicate.

### Statistics

All statistical analysis was performed using Prism 10.2.1. The individual statistical tests performed in each assay are described in the figure legends.

## DATA AVAILABILITY

The transcriptome (RNA-seq) data are deposited in NCBI’s are deposited in NCBI’s Gene Expression Omnibus (GEO) and can be accessed through GEO Series accession ID: GSE260902.

## ACKNOWLEDGEMENTS

We thank the past and present members of the Fries lab for their input in this manuscript, as well as Dr. Sian Piret who helped with performing the Seahorse Analysis. We also thank Dr. Won Hee Jung for the Sod2-GFP strain, and Dr. Tamara Doering for the mitochondrial fission/fusion mutant strains.

## AUTHOR CONTRIBUTIONS

Kyungyoon Yoo: Conceptualization, Methodology, Investigation, Visualization, Writing - Original Draft, Writing - Reviewing and Editing. Natalia Kronbauer Oliveira: Methodology, Investigation, Writing - Reviewing and Editing. Somanon Bhattacharya: Conceptualization, Methodology, Investigation, Writing - Reviewing and Editing. Bettina Fries: Supervision, Conceptualization, Resources, Writing - Reviewing and Editing, Funding Acquisition.

## SUPPLEMENTARY FIGURES AND TABLES

**Figure S1.**
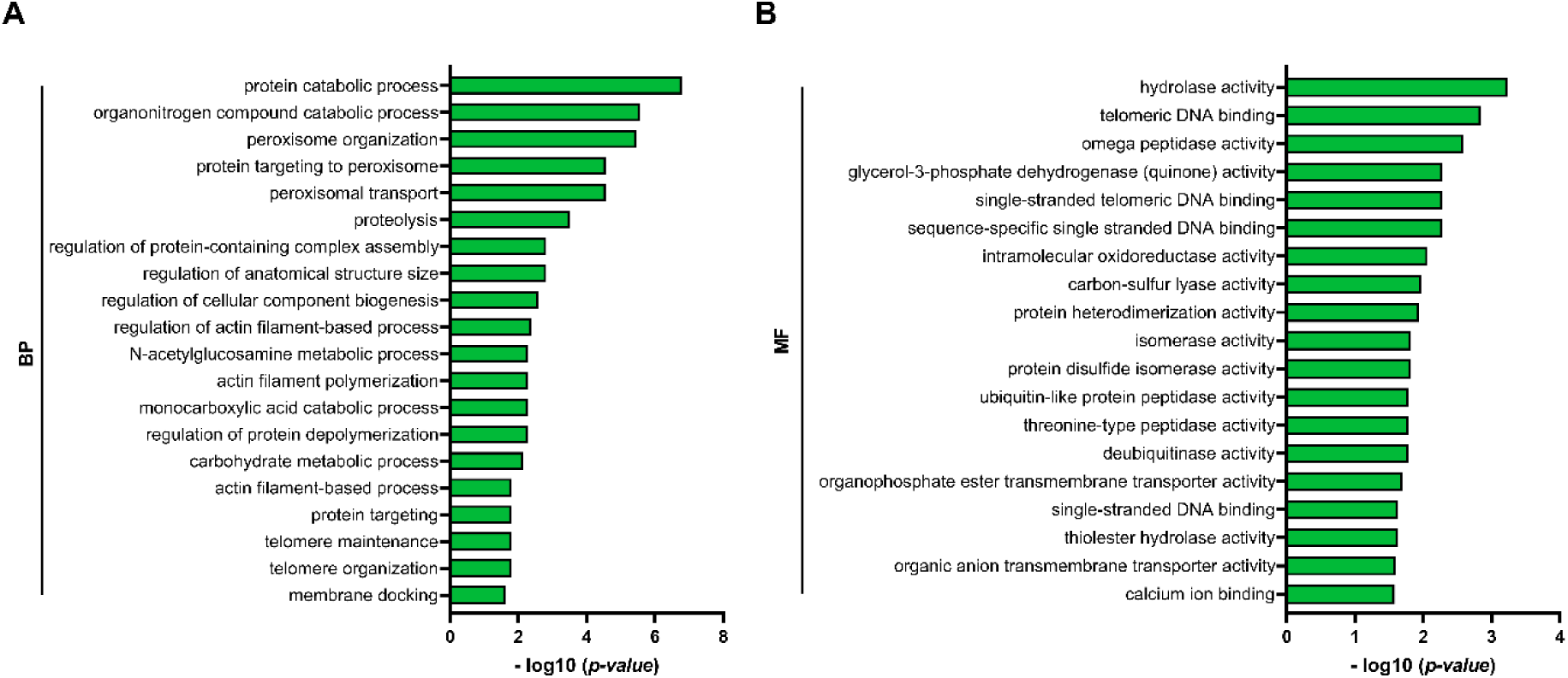
Gene Ontology Analysis of Downregulated Genes in Old *Cn* cells. Gene ontology terms of differentially expressed genes show (A) molecular function (MF) and (B) biological process (BP) that are downregulated in old *Cn* cells. Presented are the top twenty GO terms based on the lowest over-represented p-values.

**Figure S2.**
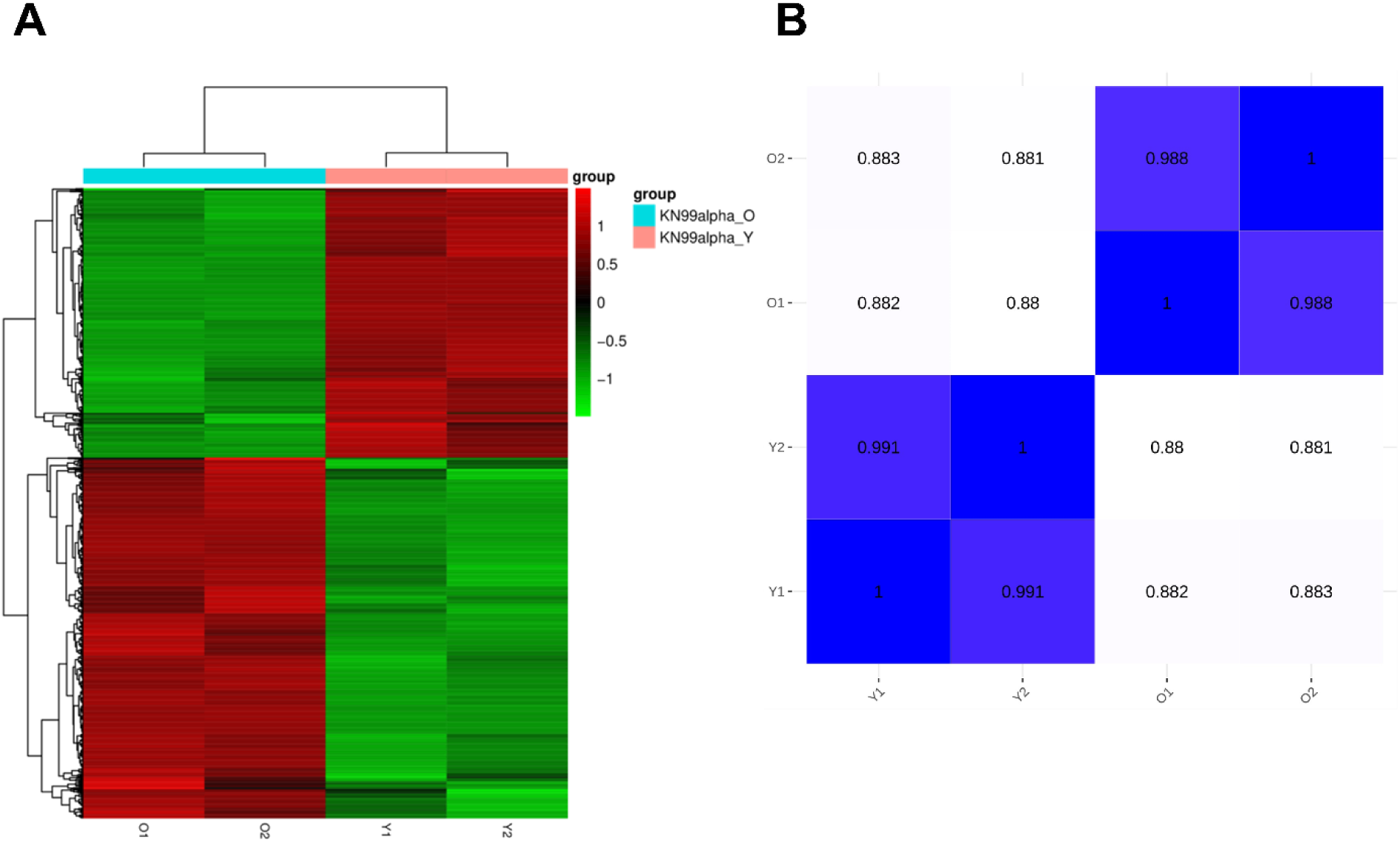
Pearson Correlation Analysis of RNA-Seq Data of Young and Old *Cn* Cells. (A) RNA-Seq heatmap showing the top all differentially expressed genes in young (KN99alpha Y) and old cells (KN99alpha O). (B) Heatmap of Pearson correlation coefficient matrix (with values presented for each pairwise comparisons) for biological replicates of young (Y1, Y2) and old (O1, O2) cells.

**Figure S3.**
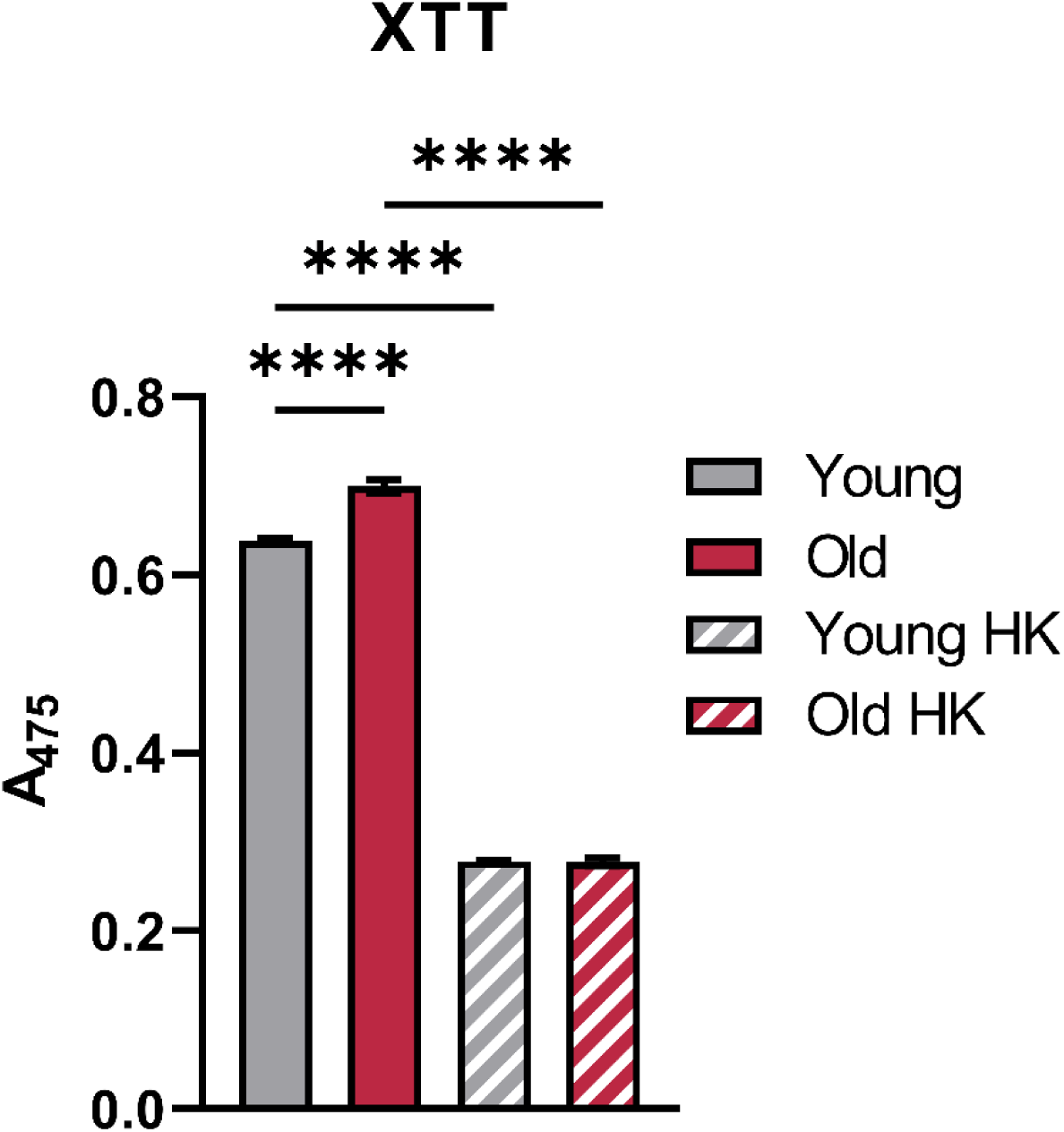
Old Cells Exhibit Higher XTT Activity. Cellular XTT [2,3-bis(2-methoxy-4-nitro-5-sulfophenyl)-2H-tetrazolium-5-carboxanilide] levels were measured using a spectrophotometer by quantifying the absorbance at 475nm. Heat-killed cells (5 mins incubation at 65°C) were used as negative controls. Bars signify the mean ± SEM of biological triplicates. One-way ANOVA with Tukey’s multiple comparisons test was used to determine the p-value. (**** p < 0.0001).

**Figure S4.**
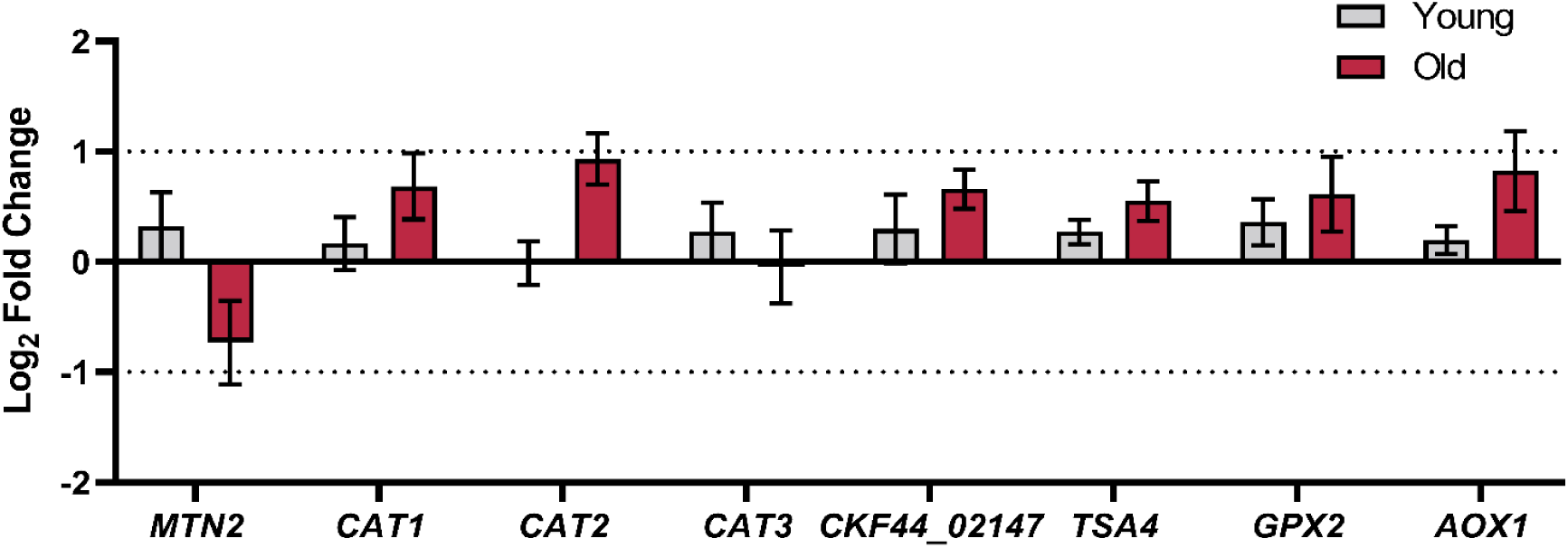
Old Cells Exhibit No Changes in Expression of Genes Encoding Some Antioxidant Genes. Some antioxidant genes were not differentially regulated between young and old *Cn* cells. mRNA expression of antioxidant genes was analyzed by RT-qPCR in young (grey bars) and old (red bars) cells. Bars signify the mean ± SEM of biological triplicates, and Student’s t-test was performed to determine the p-value (all pairwise comparisons p > 0.05). The mRNA expression is presented as Log_2_FC relative to the expression of *ACT1*, which was used as an internal control.

**Figure S5.**
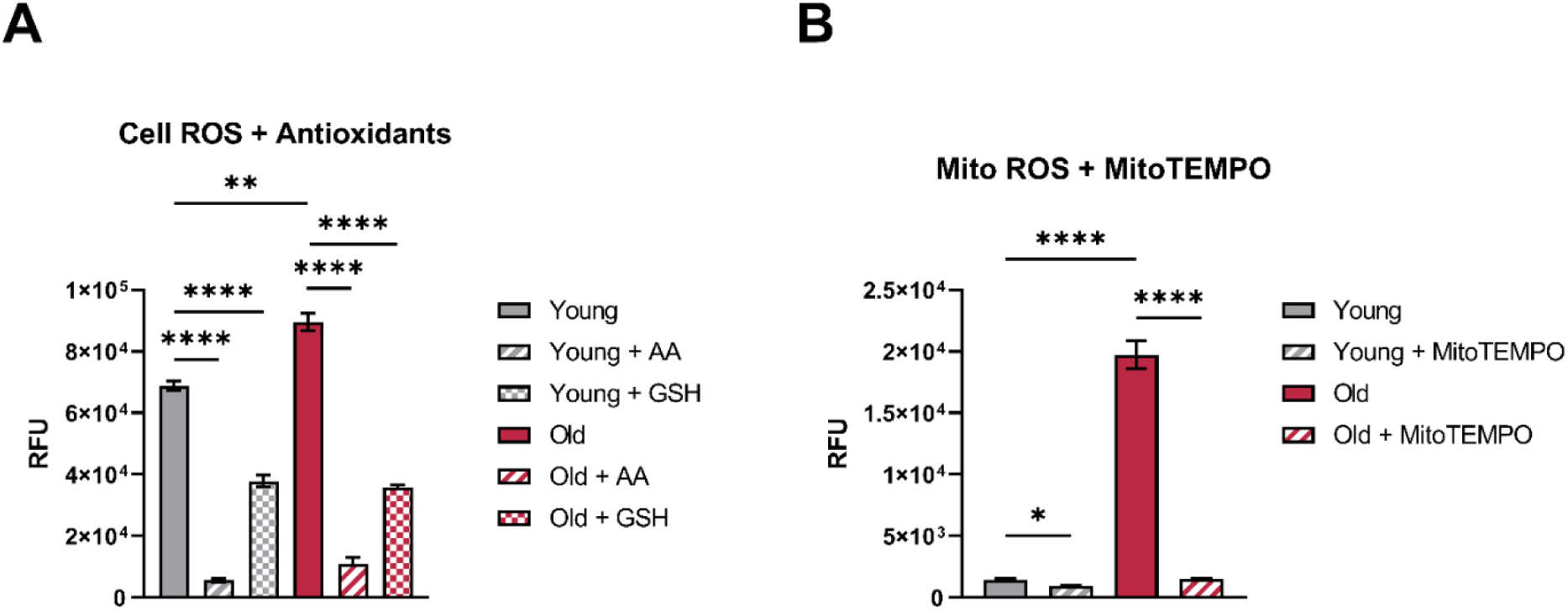
Replicative Aging in the Presence of Antioxidants Lower Reactive Oxygen Species. (A) Cellular reactive oxygen species (ROS) of young and old *Cn* cells grown in the presence of ascorbic acid (AA) and glutathione (GSH) and (B) mitochondrial ROS of young and old *Cn* cells grown in the presence of 50 µM MitoTEMPO were measured using a fluorescence plate reader by quantifying the fluorescent intensity of H_2_DCFDA and MitoSox Red respectively. Bars signify the mean ± SEM of biological triplicates. One-way ANOVA with Tukey’s multiple comparisons test was used to determine the p-value for antioxidants. (* p < 0.05, ** p < 0.01, **** p < 0.0001).

**Figure S6.**
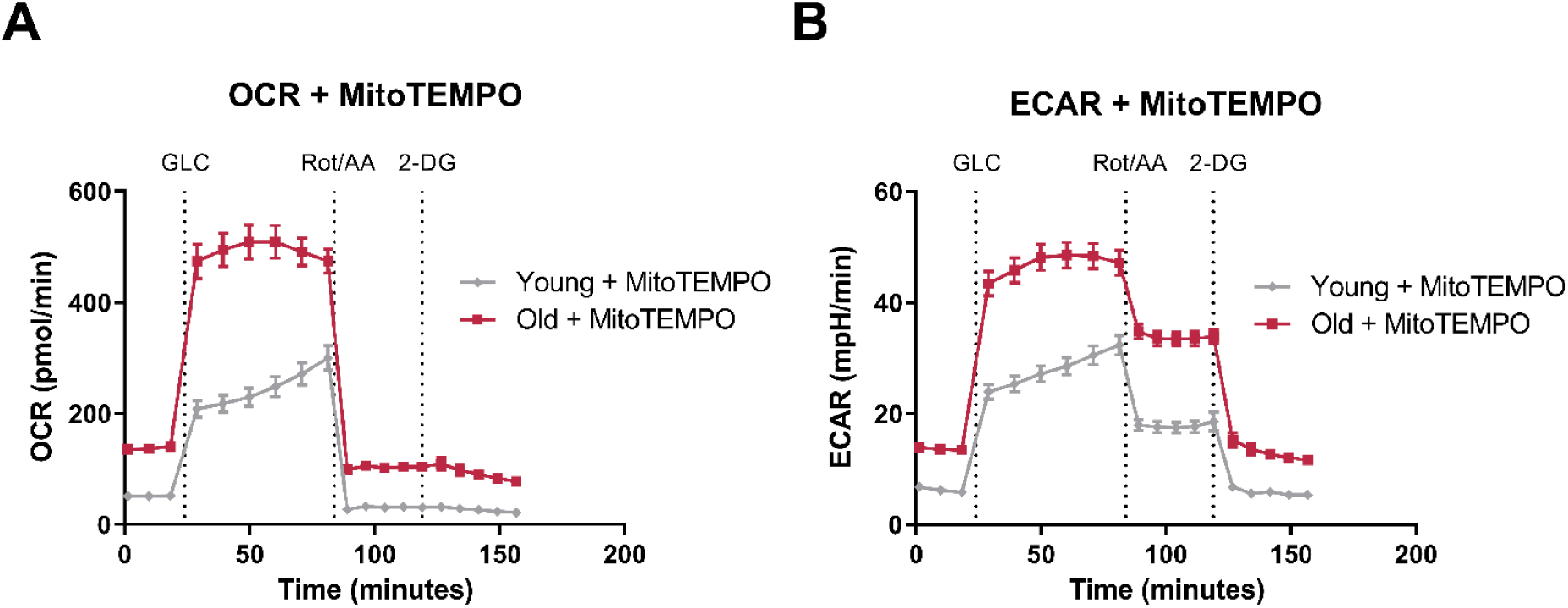
MitoTEMPO Does not Affect the Age-associated Differences in Metabolic Flux. (A) Oxygen consumption rate (OCR) and (B) extracellular acidification rate (ECAR) profiles of young and old cells were generated from the Seahorse XF glycolysis stress test following injection of glucose (GLC), Rotenone/Antimycin A (Rot/AA), and 2-deoxyglucose (2-DG). Plotted values signify the mean ± SEM of 16 replicates.

**Table S1.** Strains used in this study.

| *Cryptococcus neoformans* Strains | Source or Reference |
| --- | --- |
| KN99α | WT |
| KN99α Sod2-GFP | (70) |
| KN99α *fis1Δ* | (30) |
| KN99α *mdv1Δ* | (30) |
| KN99α *dmn1Δ* | (30) |
| KN99α *fis1Δ* | (30) |
| KN99α Fzo1<sup>OE</sup> | (30) |

**Table S2.**
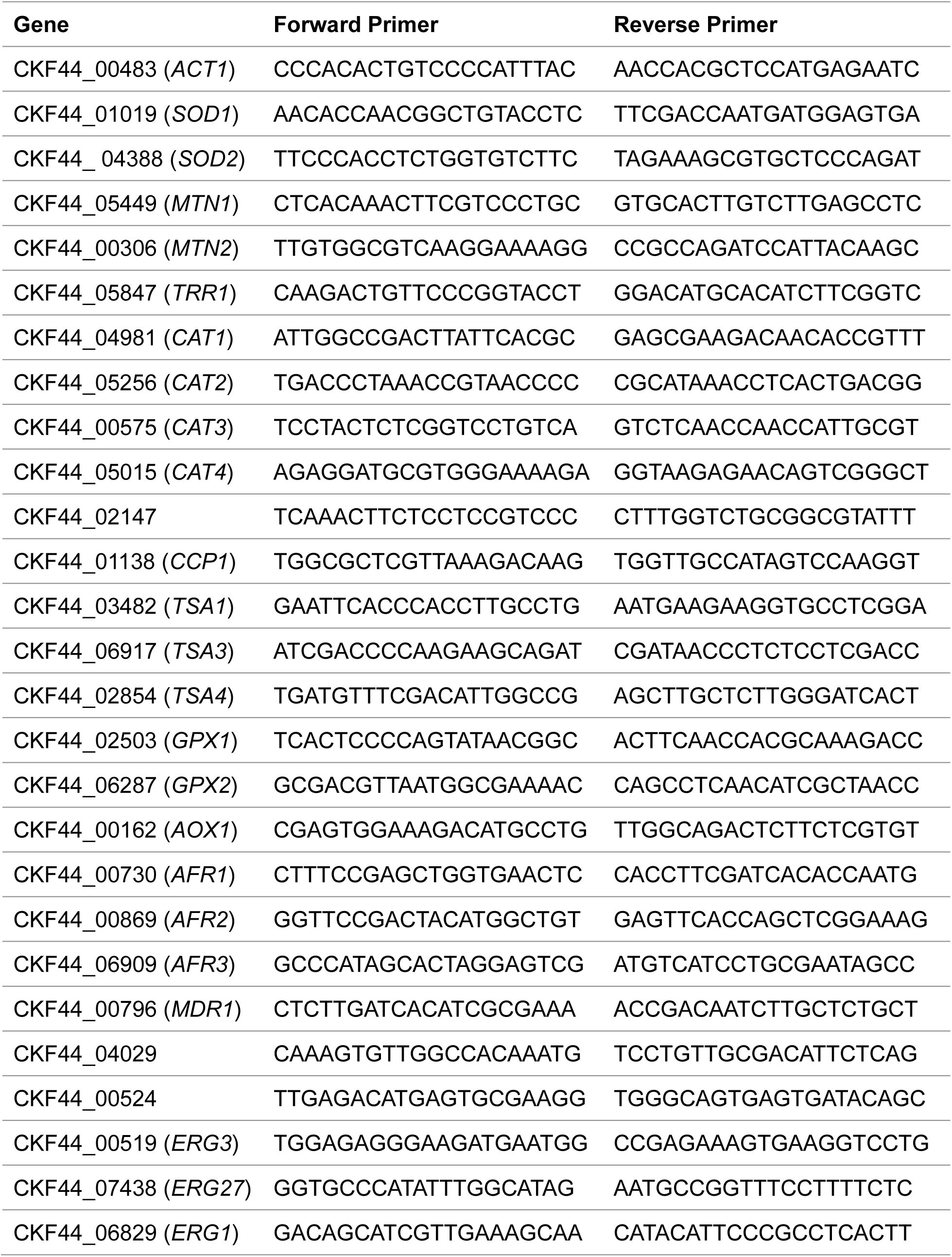
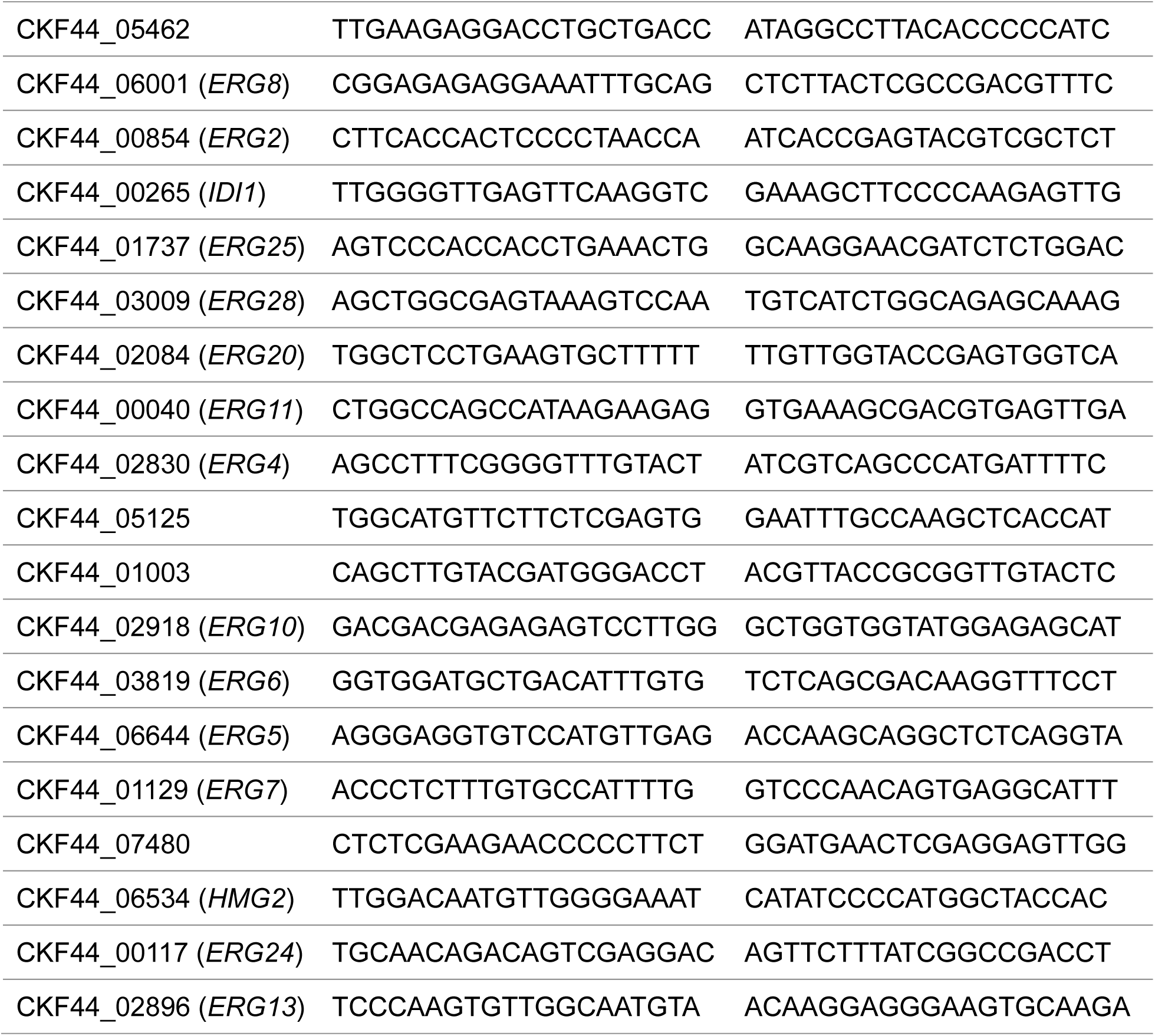
Oligonucleotides used in this study.

